# Stereochemistry Determines Immune Cellular Responses to Polylactide Implants

**DOI:** 10.1101/2022.10.27.514118

**Authors:** Chima V. Maduka, Mohammed Alhaj, Evran Ural, Maxwell M. Kuhnert, Oluwatosin M. Habeeb, Anthony L. Schilmiller, Kurt D. Hankenson, Stuart B. Goodman, Ramani Narayan, Christopher H. Contag

## Abstract

Repeating L- and D-chiral configurations determine polylactide (PLA) stereochemistry which affects its thermal and physicochemical properties, including degradation profiles. Clinically, degradation of implanted PLA biomaterials promotes prolonged inflammation and excessive fibrosis, but the role of PLA stereochemistry is unclear. Additionally, although PLA of varied stereochemistries cause differential immune responses in-vivo, this observation has yet to be effectively modeled in-vitro. A bioenergetic model was applied to study immune cellular responses to PLA containing > 99% L-lactide (PLLA), > 99% D-lactide (PDLA) and a 50/50 melt-blend of PLLA and PDLA (stereocomplex PLA). Stereocomplex PLA breakdown products increased IL-1β, TNF-α and IL-6 protein levels but not MCP-1. Expression of these proinflammatory cytokines is mechanistically driven by increases in glycolysis in primary macrophages. In contrast, PLLA and PDLA degradation products selectively increase MCP-1 protein expression. Whereas both oxidative phosphorylation and glycolysis are increased with PDLA, only oxidative phosphorylation is increased with PLLA. For each biomaterial, glycolytic inhibition reduces proinflammatory cytokines and markedly increases anti-inflammatory (IL-10) protein levels; differential metabolic changes in fibroblasts were observed. These findings provide mechanistic explanations for the diverse immune responses to PLA of different stereochemistries, and underscore the pivotal role of immunometabolism on the biocompatibility of biomaterials applied in medicine.

## 1. Introduction

The host immune response to biomaterials and material properties comprise the most important considerations for biodegradable implants to safely perform their intended function^1^. An important material property of polylactide (PLA), a biodegradable biomaterial widely used to make implants, is its stereochemistry as PLA hydrolyzes into D- or L-lactic acid; with meso-lactide being optically inactive. The stereochemistry of PLA determines its mechanical and thermal properties, crystallinity and degradation rate; higher L-lactide content increases mechanical strength, melting point and crystallinity but slows degradation^2^. Consequently, whereas PLA containing > 99% L-lactide (PLLA) takes more than 5 years to completely breakdown after implantation in animals and humans, PLA containing > 99% D-lactide (PDLA) takes only 1.5 years^1^. Furthermore, melt-blending a 50/50 ratio of PLLA and PDLA results in stereocomplex PLA that has higher mechanical strength, melting point and crystallinity, and slower degradation rate than either homopolymer because of its more compact crystal orientation^3^. Hydrolytic degradation of PLLA and stereocomplex has been characterized^4^. Degradation products of PLA, including oligomers and monomers of lactic acid, drive adverse host immune responses such as long-term inflammation and excessive fibrosis which impair PLA medical devices from performing their diagnostic, therapeutic or regenerative functions.

Historically, adverse responses to PLA degradation have been attributed to the accompanying acidity from breakdown products^5^. Recently, an alternative mechanism for immune cell activation to bulk PLA and its hydrolytic degradation products was proposed using a bioenergetic in-vitro model simulating in-vivo events^6^. In the study, altered bioenergetic homeostasis and functional metabolic profiles were discovered to underlie adverse immune responses to amorphous and crystalline PLA degradation. Higher crystallinity delayed onset of adverse immune responses, likely from faster degradation in amorphous compared to semi crystalline PLA^6^. However, spanning over decades, adverse responses to PLA have been controversial. While there have been some studies showing only mild proinflammatory responses to PLA degradation products^7–9^, other studies have demonstrated that PLA degradation is accompanied by severe adverse immune responses^10–14^ which could necessitate intervention in patients^15–17^. Differing PLA stereochemistry, which determines its diverse physicochemical and thermal properties, could account for these markedly different outcomes after surgical implantation. To test this hypothesis, breakdown products from PLLA and PDLA, which are widely applied materials in patients, as well as stereocomplex PLA were examined herein to determine the role of PLA stereochemistry in activating macrophages^18^ and fibroblasts^14^, key immune cells involved in host immune responses.

## 2. Materials and methods

### 2.1. Polylactide (PLA) materials and extraction

Polylactide (PLA) containing > 99% L-lactide (PLLA) and > 99% D-lactide (PDLA) were obtained from NatureWorks LLC as PLA L175 and PLA D120, respectively. To produce stereocomplex PLA, pre-mixtures of 50% PLLA and 50% PDLA were melt-blended in a co-rotating twin-screw extruder type ZSE 27 HP–PH (Leistritz). The screws used possessed a diameter of 27 mm and an L/ D ratio of 40/ 1; screw elements are interchangeable to allow for optimization of varied material specifications. The temperature profile range was 150 to 220 °C. After quenching the filament in a cold-water bath, the product was pelletized and then placed in a tray for drying for 24 h at 45 °C. Prior to using them, polylactide pellets were confirmed to be of similar curved surface area and sterilized by autoclaving at 121 °C for 20 minutes^19^. According to methods outlined by the International Standard Organization (ISO 10993-5:2009 - Biological evaluation of medical devices), extracts were made as previously described^6^. Briefly, we suspended 4g of biomaterial pellets, having similar surface areas, in 25 mL of complete medium (see section 2.5). After 12 days (d) at 250 rpm at 37 °C, the medium containing PLA breakdown products (called extracts) was decanted and used to treat cells for experiments. Control medium (without PLA pellets) underwent similar exposures to exclude potential confounders. Similarly, for pH testing and comparison to other studies, extraction was performed in Milli-Q water. The exact amount of extracts derived from complete medium and used to treat cells is indicated in figure legends.

### 2.2. pH measurements

pH of extracts was assessed using an Orion Star A111 Benchtop pH Meter (ThermoFisher Scientific) under room temperature conditions (20 °C).

### 2.3. Bioenergetic assessment

ATP levels in live cells were assessed by bioluminescence on the IVIS Spectrum in vivo imaging system (PerkinElmer) after adding 150 μg/mL of D-luciferin (PerkinElmer). For lysed cells, the standard ATP/ADP kits (Sigma-Aldrich) were used according to manufacturer’s instructions.

### 2.4. Microscopy

The DeltaVision imaging system and softWoRx software (GE Healthcare) were used for imaging in z-stack at 40x magnification.

### 2.5. Cells

Both mouse embryonic fibroblast cell line (NIH 3T3; ATCC) and murine primary bone-marrow derived macrophages (BMDMs) were used as previously described in bioenergetic models used in studying polylactide biocompatibility^6^. BMDMs were sourced from C57BL/6J mice (Jackson Laboratories) of 3-4 months^20,21^. In addition, NIH 3T3 cells with a Sleeping Beauty transposon plasmid (pLuBIG) having a bidirectional promoter driving an improved firefly luciferase gene (fLuc) and a fusion gene encoding a Blasticidin-resistance marker (BsdR) linked to eGFP (BGL)^22^ was used to assess ATP levels in live cells by making ATP the rate limiting factor as previously described^6^. This allowed morphological and bioenergetic changes in live cells to be concurrently monitored^23,24^. In each well of a 96-well plate, 5,000 fibroblasts or 50,000 macrophages were cultured in complete medium containing DMEM, 10% heat-inactivated Fetal Bovine Serum and 100 U/mL penicillin-streptomycin (all from ThermoFisher Scientific). Cells in each well of a 96-well plate had a total of 200 μL complete medium, out of which some was control or PLA extract (as indicated in each figure legend).

### 2.6. Materials

For pharmacologic inhibition of glycolytic flux, 3-(3-pyridinyl)-1-(4-pyridinyl)-2-propen-1-one (MilliporeSigma), 2-deoxyglucose (MilliporeSigma) and aminooxyacetic acid (Sigma-Aldrich) were used.

### 2.7. Cell viability

As previously described, cell viability was assessed using the crystal violet assay^25^. Briefly, out of a total of 200 μL of complete medium in each well of a 96-well flat-bottom plate, 150 μL was discarded. Afterwards, 150 μL of 99.9% methanol (MilliporeSigma) was added to fix cells, then removed. Similarly, 100 μL of 0.5 % crystal violet made in 25 % methanol was used to stain cells for 20 mins, after which wells were emptied. Each well was washed twice using 1x phosphate buffered saline for 2 mins, then emptied. Absorbance was acquired at 570 nm using a SpectraMax M3 Spectrophotometer (Molecular Devices) and SoftMax Pro Software (v. 7.0.2; Molecular Devices).

### 2.8. Functional metabolism

Oxygen consumption rate (OCR), extracellular acidification rate (ECAR) and lactate-linked proton efflux rate (PER) were measured using the Seahorse XFp and XFe-96 Extracellular Flux Analyzer (Agilent Technologies) as previously described^6^. After each time point, the medium in each well was washed off using Seahorse DMEM medium (pH 7.4); this removed any PLA breakdown products in contact with seeded cells. Afterwards, plates were incubated for an hour in a non-CO_2_ incubator, according to manufacturer’s instruction. Wave software (v. 2.6.1) was used to export the data as means ± standard deviation (SD).

### 2.9. Chemokine and cytokine measurements

Levels of proinflammatory and anti-inflammatory cytokines and chemokines were evaluated from BMDM supernatants using using a MILLIPLEX MAP mouse magnetic bead multiplex kit (MilliporeSigma)^26^ to assess for IL-6, MCP-1, TNF-α, IL-1β, IL-4, IL-10, IFN-λ and 1L-13 protein expression in supernatants. Using the glycolytic inhibitor, 3PO, expectedly decreased cytokine values to < 3.2 pg/ mL in some experiments. For statistical analyses, those values were expressed as 3.1 pg/ mL. Additionally, IL-6 ELISA kits (RayBiotech) for supernatants were used according to manufacturer’s instructions.

### 2.10. Optical rotation

Polarimetry was used to characterize the L-content and optical purity of the PLA samples with a P-2000 polarimeter (Jasco). The optical rotation, [*α*]_25_, was measured and averaged for three samples of each polymer in chloroform (Omnisolv), at a concentration of 1 g/100 mL. Conditions were set at 25 °C and 589 nm wavelength. Sucrose was used as a standard reference material, and its specific optical rotation was approximately 67°.

### 2.11. Gel permeation chromatography

Gel permeation chromatography (GPC) was conducted to characterize the polymer molecular weights using a 600 controller (Waters) equipped with Optilab T-rEX refractive index (RI) and TREOS II multi-angle light scattering (MALS) detectors (Wyatt Technology Corporation), and a PLgel 5μm MIXED-C column (Agilent Technologies) with chloroform eluent (1 mL/min). Polystyrene standards (Alfa Aesar) with M_n_ ranging from 35,000 to 900,000 Da were used for calibration. Dissolving stereocomplex PLA in chloroform/1,1,1,3,3,3-hexafluoro-2-propanol [90/10(v/v)] has been described for obtaining its molecular weight^27^. However, this did not dissolve our stereocomplex PLA even after heating below boiling temperatures of chloroform, likely from the high crystallinity thickness of stereocomplex PLA^27^. As such, we refer to the molecular weights of the PLLA and PDLA used in producing stereocomplex PLA.

### 2.12. Differential scanning calorimetry

Differential scanning calorimetry (DSC) was conducted with a DSC Q20 (TA Instruments) to analyse the melting temperature (T_m_), glass transition temperature (T_g_), and percent crystallinity of the PLA grades. For PLLA and PDLA, the temperature was first equilibrated to 0 °C, then ramped up to 200 °C at a heating rate of 10 °C/ min; temperature was then held isothermally for 5 minutes. Afterwards, the sample was cooled back to 0 °C at a rate of 10 °C/min, then held isothermally for 2 minutes. Finally, the material was heated back to 200 °C at 10 °C/ min. However, for stereocomplex PLA, after equilibrating to 0 °C, the temperature was ramped up to 260 °C at a heating rate of 10 °C/ min. A second heating scan was not run because, above its melting temperature (~240 °C), stereocomplex PLA thermally dissociates into its constituent homopolymers.

### 2.13. Attenuated total reflectance – Fourier transform infrared (ATR–FTIR) spectroscopy

PLA stereocomplexation was confirmed using an IRAffinity-1 spectrophotometer (Shimadzu) where the changes in the conformation of PLA chains could be observed using ATR-FTIR spectroscopy as previously described^28^.

### 2.14. Liquid Chromatography-Electrospray ionization mass spectrometry (LC-ESI-MS

Extracts of PLLA, PDLA and stereocomplex PLA made in Milli-Q water were directly analyzed by LC-MS using a Thermo Q-Exactive mass spectrometer interfaced with a Thermo Vanquish UHPLC. 5 uL of extract was injected onto a Waters Acquity BEH-C18 UPLC column (2.1×100 mm) and lactic acid oligomers were separated using the following gradient: initial conditions were 98% mobile phase A (0.1% formic acid in water) and 2% mobile phase B (acetonitrile + 0.1% formic acid), hold at 2% B until 1.0 min, linear ramp to 99% B at 7.0 min and hold at 99% B until 8 min, return to 2% B at 8.1 min and hold at 2% B until 10 min. The flow rate was 0.3 ml/min and the column temperature was 40°C. Ions were generated by electrospray ionization in negative mode with a capillary voltage of −2.5 kV and source gas flow and temperature settings were set as the source auto-defaults for an LC flow rate of 0.3 ml/min. MS and MS/MS data were acquired using a data-dependent MS method with survey scans acquired at 70,000 resolution (scan range m/z 80-1200) and MS/MS scans for the top 5 ions acquired at 17,500 resolution with an isolation width of 1.0 m/z and stepped normalized collision energies settings of 10, 30 and 60.

### 2.15. Statistics and reproducibility

Data were presented as mean with standard deviation (SD). For data analysis, statistical software (GraphPad Prism) was used with significance level set at p < 0.05. Specific details of statistical tests and sample sizes are provided in figure legends.

## 3. Results

We validated the physicochemical and thermal properties of polylactide (PLA) containing > 99% L-lactide (PLLA), > 99% D-lactide (PDLA) and stereocomplex PLA (melt-blend of 50/50 PLLA and PDLA) prior to using them (Table S1). Stereocomplexation was confirmed by both differential scanning calorimetry (DSC) thermograms and attenuated total reflectance–Fourier transform infrared (ATR–FTIR) spectroscopy. DSC thermograms revealed a melting peak of 240 °C (Fig. S1A). With ATR-FTIR spectroscopy, the α helix (wavenumber 921 cm^−1^) which is characteristic of PLLA and PDLA is transformed into a more compact β helix (wavenumber 908 cm−1) in stereocomplex PLA (Fig. S1B, C)^28^. Using a bioenergetic model we had developed and optimized^6^, we degraded the different types of PLA to obtain breakdown products over 12 days (d), also referred to as extracts in complete (serum containing) medium. There were no changes in pH over the 12 d extraction period for serum-containing control (no polylactide) medium (pH = 8.0), PLLA (pH = 8.0), PDLA (pH = 8.1) and stereocomplex PLA (pH = 8.1) extracts used on cells. In contrast, extraction in water resulted in changes between control (no polylactide; pH = 8.2), PLLA (pH = 7.5), PDLA (pH = 7.7) and stereocomplex PLA (pH = 7.3) extracts; this decrease in pH with water-derived stereocomplex PLA extract has been reported^4^.

We had previously quantified monomeric D- and L-lactic acid content in PLA extracts after 12 d of extraction^6^. To evaluate oligomers of lactic acid released in different extracts, we used liquid chromatography coupled with electrospray ionization-mass spectrometry (LC-ESI-MS)^4^. Extracts derived from equal amounts of polymer were analyzed in negative ion mode where the expected mass to charge ratio (m/z) of lactic acid is 89.024418^29^, with higher order polylactic acid oligomers having additional C_3_H_4_O_2_ units with a mass of 72.021^4^. Therefore, the expected series was 89.024418 (a monomeric lactic acid unit), 161.045547 (2 units), 233.066676 (3 units), 305.087806 (4 units), 377.108935 (5 units), 449.130064 (6 units), 521.150097 (7 units),593.172323 (8 units), and so on. The PDLA extracts showed the highest abundance of oligomeric lactic acid released (Fig. S2). Oligomers up to eight lactic acid units were detectable in PDLA samples with free lactic acid being the most abundant (Fig. S2). The stereocomplex PLA samples had oligomers up to seven units detected, but at levels approximately 1-12% of that for the PDLA samples (Fig. S2). PLLA samples had the lowest levels of detectable lactic acid oligomers (only up to six detectable units) with levels ~0.5-5% of the PDLA samples (Fig. S2).

After exposure to PLLA, PDLA and stereocomplex PLA extracts, there were no alterations in ATP levels over time in live (Fig. 1A) or lysed (Fig. 1B) fibroblasts in comparison to untreated cells, despite tendencies for changes on days 3 and 7. In addition, there were no changes in microscopic appearance of fibroblasts exposed to PLLA or PDLA extracts compared to untreated cells (Fig. 1C). However, fibroblasts exposed to stereocomplex PLA extract tended to be more rounded than stellar in appearance (Fig. 1C). In contrast to fibroblasts, primary bone marrow-derived macrophages exposed to PLLA, PDLA or stereocomplex PLA extracts expressed higher ATP levels than untreated cells on days 7 and 11 (Fig. 2). Evaluation of dose-bioenergetic response by adding different volumes of polylactide extracts revealed changes in ATP levels for all doses (Fig. S3), and this guided our subsequent studies.

**Fig. 1.**
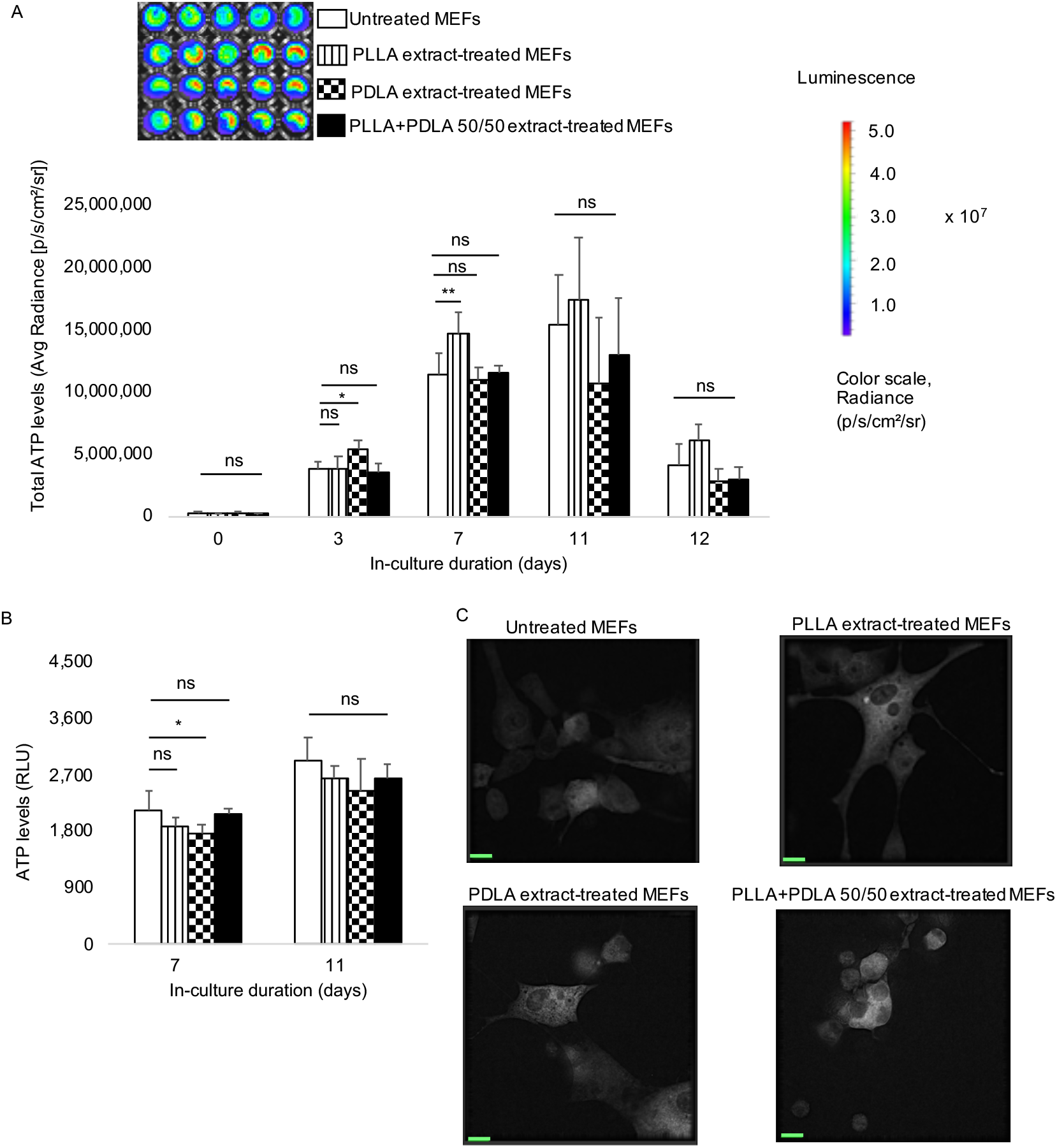
A) Bioenergetic (ATP) levels in mouse embryonic fibroblasts (MEFs) are mostly unaltered after exposure to breakdown products (extracts) of polylactide containing >99% L-isomer (PLLA), >99% D-isomer (PDLA) or a 50/50 melt-blend of PLLA and PDLA (stereocomplex PLA) over time in live cells. Representative images for day 7 are shown. B) In lysed cells, ATP levels are mostly unchanged following exposure to PLLA, PDLA and stereocomplex PLA extracts. C) There were no apparent microscopic changes in MEFs exposed to PLLA or PDLA; cells exposed to stereocomplex PLA extract appeared more rounded in comparison to untreated cells on day 7 (scale bar, 15 μm). Not significant (ns), *p<0.05, **p<0.01, mean (SD), n=3-5, one-way ANOVA followed by Tukey’s post-hoc test; 150 μl of control or extract was used.

**Fig. 2.**
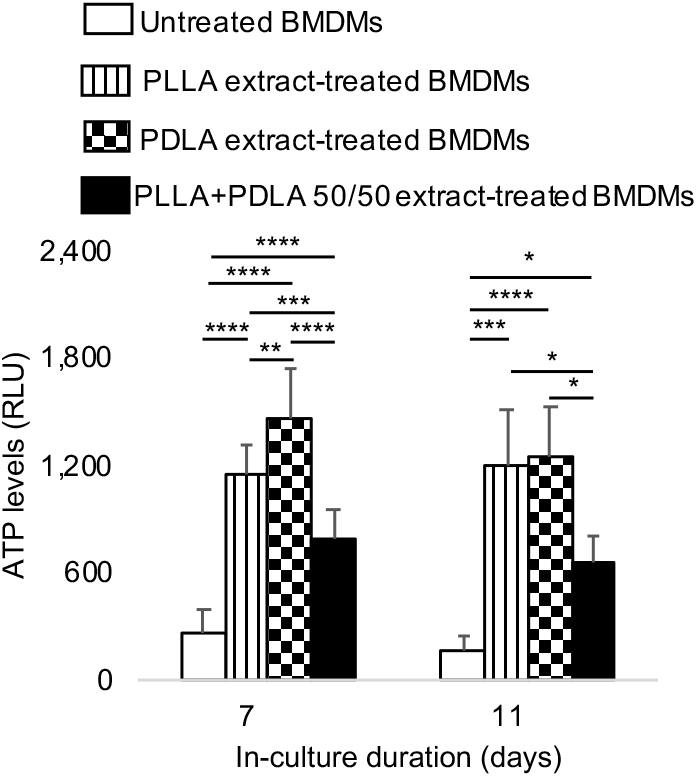
In primary bone marrow-derived macrophages (BMDMs), bioenergetics is altered after prolonged exposure to breakdown products (extracts) of polylactide containing >99% L-isomer (PLLA), >99% D-isomer (PDLA) or a 50/50 melt-blend of PLLA and PDLA (stereocomplex PLA) over time. *p<0.05, **p<0.01, ***p<0.001, ****p<0.0001, mean (SD), n=4-10, one-way ANOVA followed by Tukey’s post-hoc test; 150 μl of control or extract was used.

Next, we sought to find out whether bioenergetic changes were affected by cell numbers. Using the crystal violet assay^25^, we observed that numbers of fibroblasts were similar after exposure to PLLA, PDLA or stereocomplex PLA extract in comparison to untreated cells over time (Fig. S4). Observed changes in numbers of macrophages (Fig. S5) were not a confounder after normalizing ATP levels in macrophages (Fig. S6). Bioenergetic alterations from amorphous and crystalline polylactide degradation could be the result of changes in oxidative phosphorylation and glycolytic flux in immune cells^6^. To determine whether PLLA, PDLA or stereocomplex PLA degradation alters functional metabolism, we used the Seahorse assay to measure oxygen consumption rate (OCR; a measure of oxidative phosphorylation), extracellular acidification rate (ECAR; a measure of glycolytic flux) and proton efflux rate (PER; a measure of monocarboxylate transporter function, where monocarboxylate transporters are responsible for the bidirectional movement of proton linked lactate across cell membranes)^30–32^.

Compared to untreated macrophages, extracts of PLLA and PDLA increased OCR (Fig. 3A, B). However, similar amounts of stereocomplex PLA extract did not affect OCR (Fig. 3C). Whereas exposure of macrophages to PLLA extract did not affect ECAR (Fig. 3D), extracts of PDLA and stereocomplex PLA increased ECAR (Fig. 3E, F). Similarly, PER was unchanged after exposure to PLLA extract (Fig. 3G) but exposure to PDLA and stereocomplex PLA extracts increased PER (Fig. 3H, I), suggesting a trend for both PDLA and stereocomplex PLA which each contain ≥ 50% D-lactide.

**Fig. 3.**
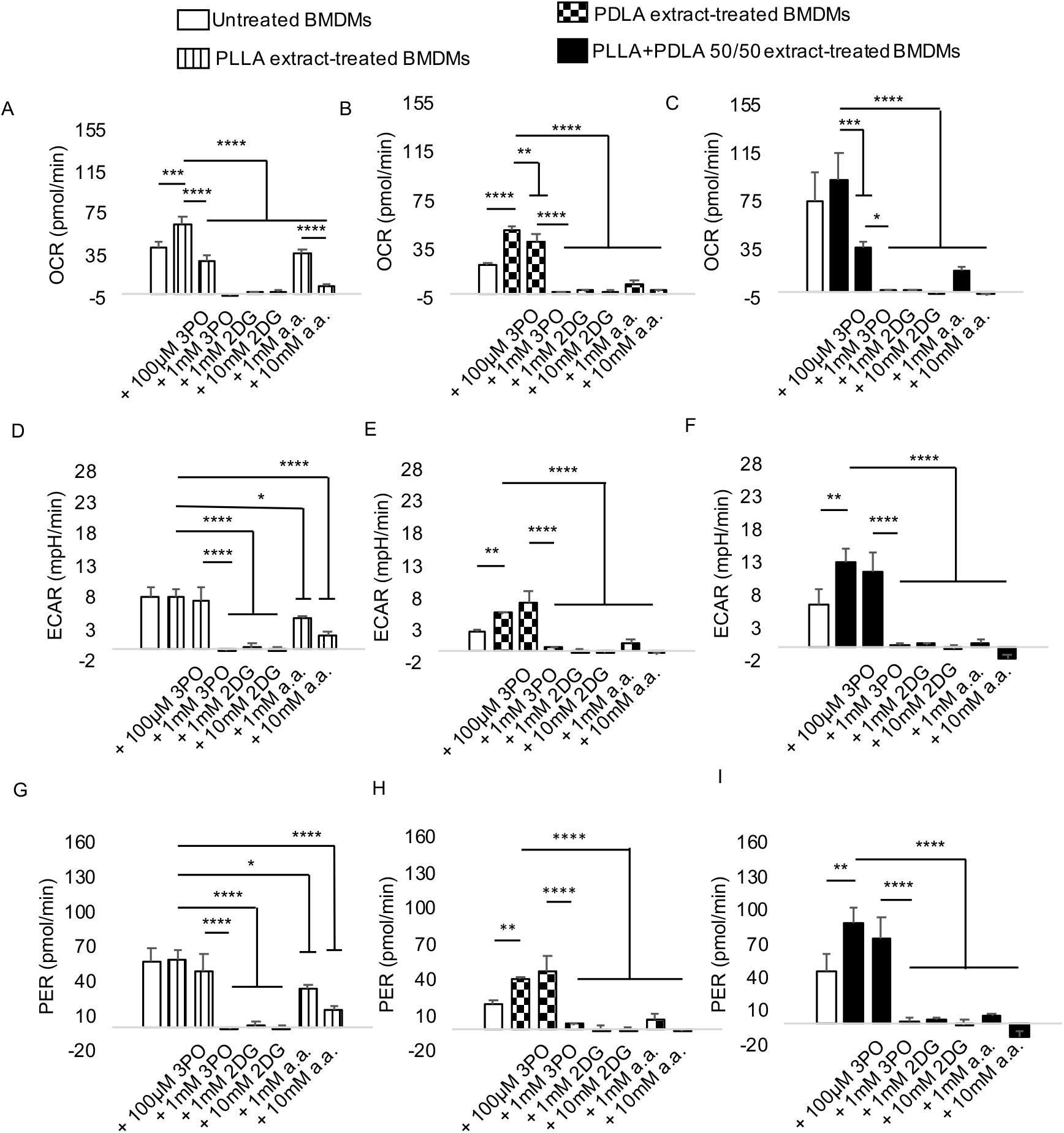
A-C) In comparison to untreated cells, oxygen consumption rate (OCR) is increased in primary bone marrow-derived macrophages (BMDMs) exposed to breakdown products (extracts) of polylactide containing >99% L-isomer (PLLA) or >99% D-isomer (PDLA) but not a 50/50 melt-blend of PLLA and PDLA (stereocomplex PLA). In cells exposed to polylactide extracts, OCR is dose-dependently controlled by addition of glycolytic inhibitors. D-I) Extracellular acidification rate (ECAR; D-F) and proton efflux rate (PER; G-I) are increased in BMDMs treated with PDLA or stereocomplex PLA but not PLLA extracts. Both ECAR and PER are controlled by addition of glycolytic inhibitors in a dose-dependent manner. *p<0.05, **p<0.01, ***p<0.001, ****p<0.0001, mean (SD), n=3, one-way ANOVA followed by Tukey’s post-hoc test; 3-(3-pyridinyl)-1-(4-pyridinyl)-2-propen-1-one (3PO), 2-deoxyglucose (2DG) and aminooxyacetic acid (a.a.); 100 μl of control or extract was used for 7 days.

For the volume of extract used in Seahorse assays, cell numbers were similar between untreated macrophages and macrophages exposed to PLLA, PDLA or stereocomplex PLA extract (Fig. S7A-C).

Lactate, the final substrate in glycolysis, is converted to pyruvate which feeds oxidative phosphorylation in the tricarboxylic acid cycle. Reasoning that modulation of glycolysis will also modulate oxidative phosphorylation, different steps in the glycolytic pathway were targeted. Three small molecule inhibitors used were 3-(3-pyridinyl)-1-(4-pyridinyl)-2-propen-1-one (3PO), 2-deoxyglucose (2DG) and aminooxyacetic acid (a.a.) which inhibit 6-phosphofructo-2-kinase/ fructose-2,6-bisphosphatase isozyme 3 (PFKFB3), hexokinase and uptake of glycolytic substrates, respectively^33–35^. Following exposure of macrophages to PLLA, PDLA or stereocomplex PLA extracts, addition of 3PO, 2DG or a.a. resulted in a dose-dependent decrease in OCR, ECAR and PER (Fig. 3A-I). Macrophages with altered metabolic profiles from exposure to PLLA, PDLA or stereocomplex PLA extracts had mildly decreased cell numbers after treatment with pharmacologic inhibitors (Fig. S7A-C).

In fibroblasts, exposure to stereocomplex PLA but not PLLA or PDLA extract increased OCR (Fig. 4A). ECAR and PER increased in fibroblasts exposed to PLLA or stereocomplex PLA extract compared to untreated cells (Fig. 4B, C). In contrast, ECAR and PER were similar in untreated fibroblasts and cells exposed to PDLA extract (Fig. 4B, C). Fibroblasts exposed to PLLA, PDLA, or stereocomplex PLA extracts expressed lower bioenergetic levels after addition of 3PO, 2DG or a.a. in a dose-dependent manner (Fig. 4D-F; Fig. S8A-C).

**Fig. 4.**
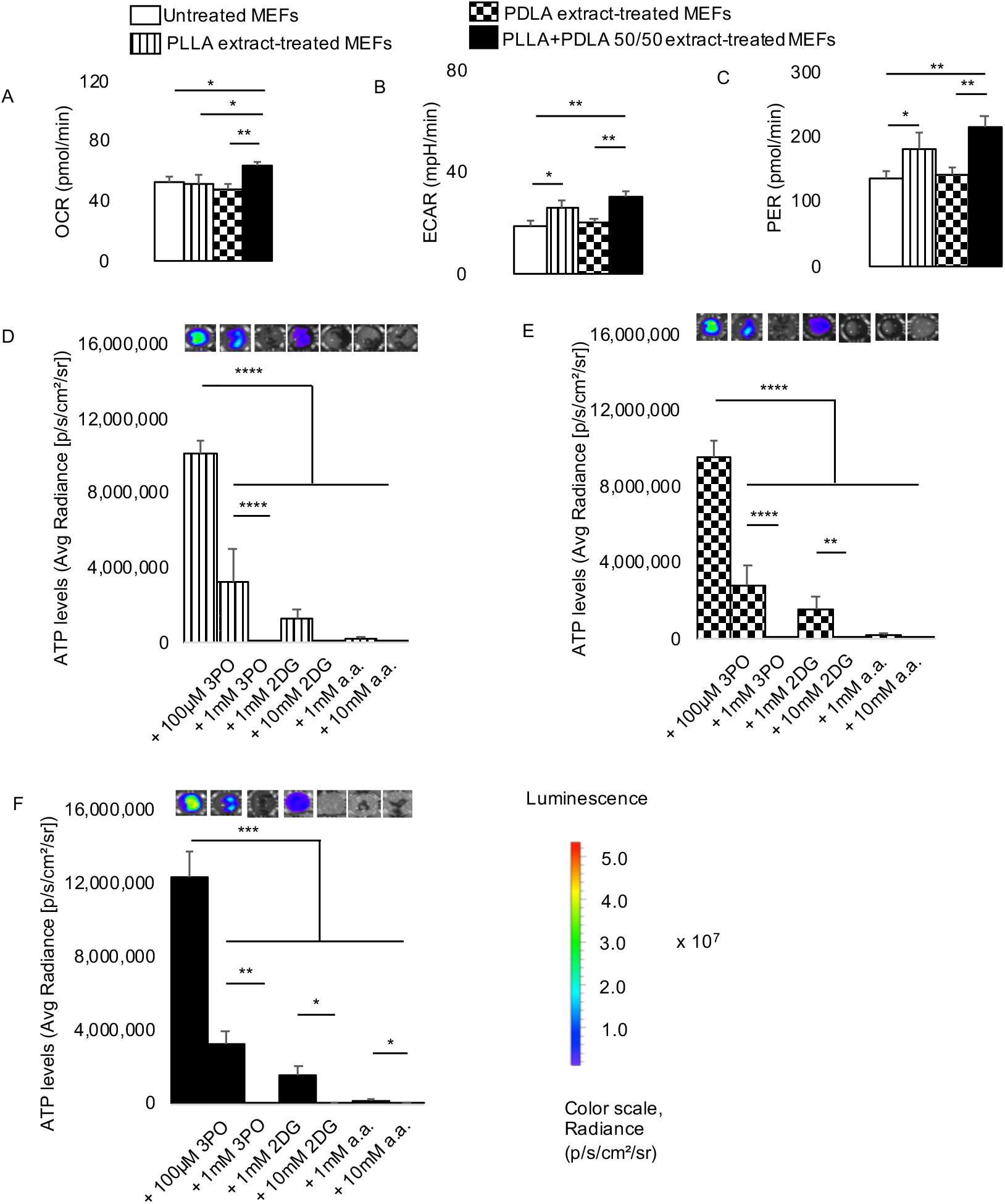
A) Oxygen consumption rate (OCR) is elevated in mouse embryonic fibroblasts (MEFs) exposed to breakdown products (extracts) of stereocomplex PLA compared to untreated, PLLA or PDLA groups. B-C) Extracellular acidification rate (ECAR; B) and proton efflux rate (PER; C) are increased in MEFs exposed to stereocomplex PLA or PLLA extracts in comparison to untreated cells. D-F) Bioenergetics is modulated in MEFs exposed to stereocomplex PLA, PLLA or PDLA extracts in a dose-dependent manner by pharmacologic inhibitors of glycolysis (representative wells are shown). *p<0.05, **p<0.01, ***p<0.001, ****p<0.0001, mean (SD), n=3-5, one-way ANOVA followed by Tukey’s post-hoc test or Brown-Forsythe and Welch ANOVA followed by Dunnett multiple comparison test; 3-(3-pyridinyl)-1-(4-pyridinyl)-2-propen-1-one (3PO), 2-deoxyglucose (2DG) and aminooxyacetic acid (a.a.); polylactide containing >99% L-isomer (PLLA), >99% D-isomer (PDLA) and a 50/50 melt-blend of PLLA and PDLA (stereocomplex PLA); 150 μl of control or extract was used for 7 days in Figure 4A-C; 100 μl of control or extract was used for 7 days in in Figure 4D-F.

To determine whether immune activation is the result of altered bioenergetics and metabolic reprogramming, we assayed levels of cytokine and chemokine expression using a magnetic bead-based technique. Both proinflammatory (MCP-1, IL-1β, TNF-α, IL-6 and IFN-γ) and anti-inflammatory (IL-4, IL-13 and IL-10) protein levels were assessed. In comparison to untreated macrophages, PLLA and PDLA but not stereocomplex PLA extract increased MCP-1 protein levels (Fig. 5A). Targeting glycolytic flux using 3PO, 2DG or a.a. consistently decreased MCP-1 expression (Fig. 5A).

**Fig. 5.**
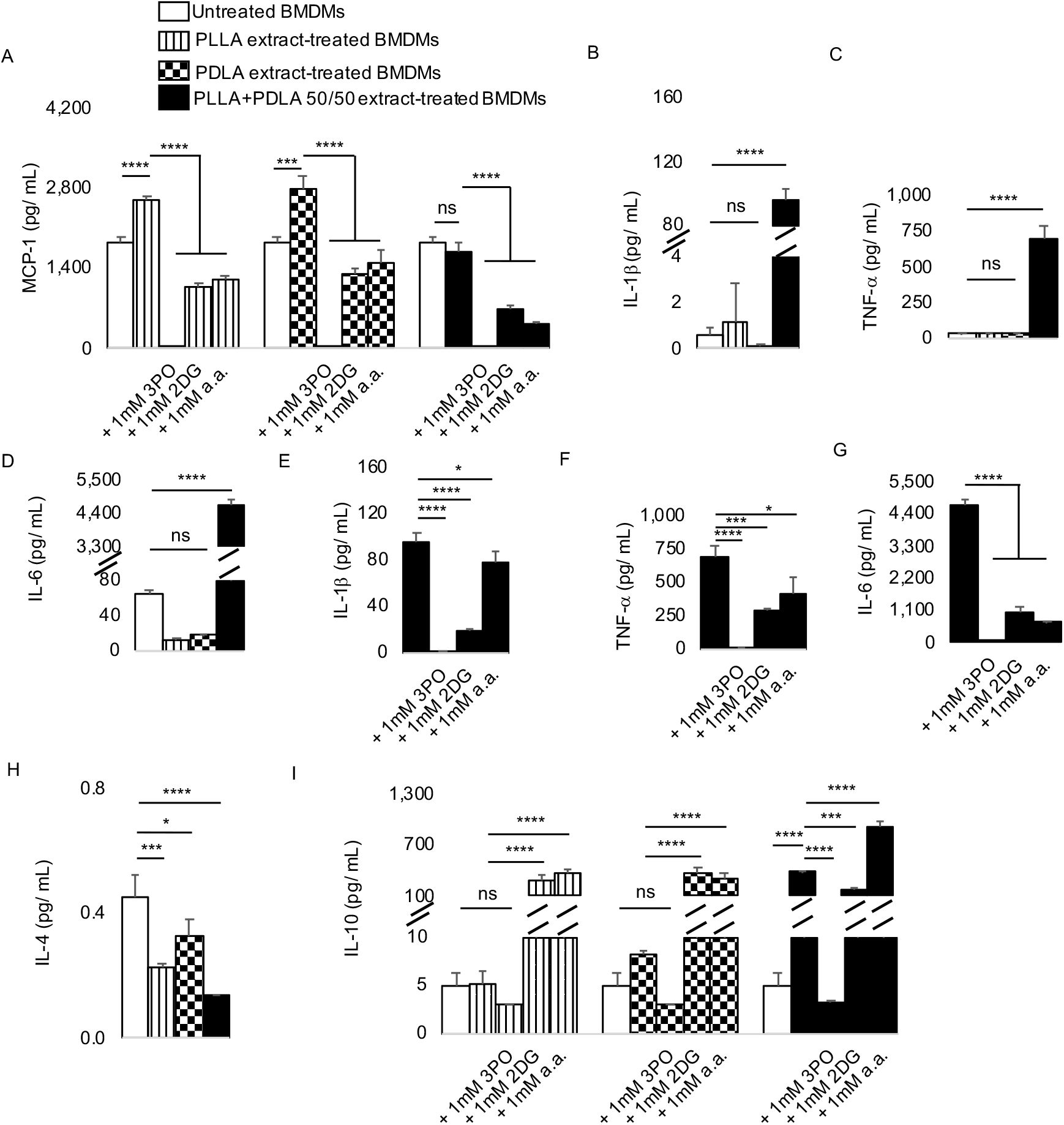
A) Compared to untreated primary bone marrow-derived macrophages (BMDMs), MCP-1 protein expression is increased in cells exposed to breakdown products (extracts) of polylactide containing >99% L-isomer (PLLA) or >99% D-isomer (PDLA) but not a 50/50 melt-blend of PLLA and PDLA (stereocomplex PLA); increased MCP-1 levels are inhibited by different glycolytic inhibitors. B-D) Whereas IL-1β, TNF-α and IL-6 protein levels are similar in BMDMs exposed to PLLA and PDLA, they are elevated in cells exposed to stereocomplex PLA extracts in comparison to untreated cells. E-G) Increased IL-1β, TNF-α and IL-6 protein expression in BMDMs exposed to stereocomplex PLA extracts is inhibited by different glycolytic inhibitors. H) IL-4 protein expression is decreased in BMDMs exposed to PLLA, PDLA or Stereocomplex PLA compared to untreated cells. I) IL-10 protein expression is unchanged in BMDMs exposed to PLLA or PDLA but not stereocomplex PLA compared to untreated cells; addition of 2-deoxyglucose (2DG) and aminooxyacetic acid (a.a.) but not 3-(3-pyridinyl)-1-(4-pyridinyl)-2-propen-1-one (3PO) increases IL-10 protein expression in comparison to cells exposed to respective polylactide extracts. Not significant (ns), *p<0.05, ***p<0.001, ****p<0.0001, mean (SD), n=3, one-way ANOVA followed by Tukey’s post-hoc test; 150 μl of control or extract was used for 7 days.

Exposure of macrophages to PLLA or PDLA extracts did not increase IL-1β (Fig. 5B), TNF-α (Fig. 5C), or IL-6 protein levels (Fig. 5D). However, similar amounts of stereocomplex PLA extract increased IL-1β, TNF-α and IL-6 levels by 175.2-, 19.1-, and 70.9-fold respectively (Fig. 5B-D). Independent studies using a different technique (ELISA) revealed similar trends for IL-6 (Fig. S9). The increased levels of IL-1β, TNF-α and IL-6 by stereocomplex PLA were decreased by targeting glycolytic flux using 3PO, 2DG or a.a. (Fig. 5E-G). There were no changes in levels of IL-13 or IFN-γ (data not shown). Macrophages exposed to PLLA, PDLA or stereocomplex PLA extracts decreased IL-4 protein expression (Fig. 5H). Interestingly, neither PLLA nor PDLA extract increased IL-10 levels (Fig. 5I). However, addition of 2DG and a.a. increased IL-10 levels by 55.8- and 71.9-fold, respectively, in macrophages exposed to PLLA extract; by 46.3- and 37-fold, respectively, in macrophages exposed to PDLA extract (Fig. 5I). Stereocomplex PLA extract increased IL-10 levels compared to untreated macrophages (Fig. 5I). Yet, inclusion of a.a. to stereocomplex PLA extract increased IL-10 levels by 2.4-fold in macrophages compared to stereocomplex PLA alone (Fig. 5I).

## 4. Discussion

There have been numerous in-vivo studies characterizing the host immune cellular response to polylactide (PLA)^7–17^. While being highly informative, the complexity of the in-vivo microenvironment where PLA is implanted could preclude a thorough understanding of mechanistic events therein. To deconvolute in-vivo events, several in-vitro studies have been undertaken^36–40^. However, many prior in-vitro models on PLA biocompatibility focused on acidity, based on prior correlation between reduced pH and the adverse immune responses elicited by PLA^5,19^. Instead, this study builds on observed alterations in bioenergetics as well as metabolic reprogramming as drivers of adverse immune responses to polyethylene wear particles^41,42^ after total joint replacements and PLA biomaterials^6^. By subcutaneously implanting sterile amorphous PLA, we validated this bioenergetic model by demonstrating metabolic reprogramming, in-vivo. We showed enhanced radiolabeled glucose (fluorodeoxyglucose; FDG) uptake in the PLA microenvironment which was abrogated by inhibiting glycolysis^6^. In agreement, persistent foreign body (sterile inflammatory) reactions to PLA devices employed after human surgeries manifest elevated FDG uptake, which could be mistaken for cancer recurrence^43,44^. By comparing glycolytic inhibition to neutralization strategies, we showed that the former was more effective in reducing CD86 (proinflammatory) expression in the amorphous PLA microenvironment, without affecting CD11b or F4/80 (macrophage recruitment) levels, supporting our in-vitro model^6^. Additionally, our bioenergetic model more robustly simulated sterile inflammatory protein expression, without the need to include IFN-γ^45^ or bacterial endotoxins^46^ in PLA in-vitro studies. Moreover, using primary macrophages directly isolated from the murine bone marrow better simulated in-vivo scenarios than monocyte-macrophage cell lines, in our model.

Bioenergetics and metabolic reprogramming are emerging mechanisms in immune cell activation^20,30,33,47,48^. Perhaps the greatest advantage of PLA-based implants is their biodegradability; it is thought that PLA degradation products are “normally” metabolized by the tricarboxylic acid cycle (TCA) to make ATP^49^. However, this had not been previously investigated. Here, we reveal that bioenergetic imbalances occur in primary macrophages following exposure to PLA containing > 99% L-lactide (PLLA), > 99% D-lactide (PDLA) or stereocomplex PLA (melt-blend of 50/50 PLLA and PDLA). This is consistent with our prior studies using amorphous PLA (with 10% D-lactide) and semi-crystalline PLA (with 1% D-lactide)^6^. Comparatively, highly crystalline PLLA and PDLA are more often used in patients because they appear to less immunogenic than other PLA types^1^. Principally composed of D- or L-lactic acid, PLLA and PDLA present a unique opportunity to investigate the role of chirality in immunological response to PLA implants.

Increased glycolytic flux, oxidative phosphorylation (OXPHOS) and monocarboxylate transporter (MCT) function are mechanistic drivers of the inflammatory response mediated by macrophages^31,47,50,51^. Therefore, our observation that PLLA, PDLA and stereocomplex PLA degradation products differentially reprogram metabolism (glycolytic flux and oxidative phosphorylation) in macrophages is exciting as it offers mechanistic insight into the different host immune responses to PLA of varied stereochemistries. Changing stereochemistries, in turn, affects mechanical and thermal properties, crystallinity and degradation rates.

Elevated glycolytic flux, OXPHOS or MCT function from exposure to PLLA, PDLA or stereocomplex PLA extracts were decreased upon addition of small molecule inhibitors, including 3-(3-pyridinyl)-1-(4-pyridinyl)-2-propen-1-one (3PO), 2-deoxyglucose (2DG) and aminooxyacetic acid (a.a.), which target different steps in glycolysis; these decreases were accompanied by slight reductions in cell numbers. However, cell numbers alone could not fully account for observed reductions in glycolytic flux, OXPHOS or MCT function. In contrast, untreated macrophages exposed to these pharmacologic inhibitors for the same duration neither exhibited decreased functional metabolism nor reduced cell numbers, suggesting selectivity for macrophages having altered metabolism only following exposure to different types of PLA^6^. Of note, one of the small molecule inhibitors, aminooxyacetic acid (a.a.; pKa: 3.16), is a stronger acid than lactic acid (pKa: 3.78). Yet, a.a. modulated inflammatory responses to stereocomplex PLA breakdown products, arguing against acidity being the sole driver of adverse immune responses as PLA degrades. Similar to macrophages, fibroblasts respond differently to degradation products of PLLA, PDLA or stereocomplex PLA in terms of bioenergetics and functional metabolism. In both macrophages and fibroblasts, there is efficient cellular uptake of 3PO, 2DG or a.a. as candidate pharmacologic agents to modulate metabolic reprogramming and altered bioenergetics.

PLA degradation products are known to drive immune cellular activation ^7–17^. Since PLLA and PDLA degrade faster than stereocomplex PLA^4^, we expected PLLA and PDLA degradation products to elicit more adverse immune cellular responses. Paradoxically, among proinflammatory cytokines, IL-1β, TNF-α and IL-6 protein levels were unchanged after exposure to PLLA or PDLA degradation products; however, stereocomplex PLA products markedly increased IL-1β, TNF-α and IL-6 protein levels. The molecular weights of PLLA and PDLA used in this study approximated each other; therefore molecular weight was not a confounder as has been shown where greater (up to ten) fold change differentiate various biomaterials that were studied^52–54^. Moreover, PLLA, PDLA and stereocomplex PLA are all highly crystalline formulations, excluding crystallinity as a confounder, since amorphous PLA degrades faster than crystalline PLA.

Extracts of PLLA and stereocomplex PLA, made in water at 37°C over durations relevant to and longer than our study’s, have been characterized. Providing critical insight, they indicate that oligomeric degradation products are observed after 13 to 19 weeks^4^. However, their PLA extraction was performed under stationary conditions. We used dynamic conditions that accelerate degradation^55^. By comparing to PLLA, they demonstrated that shorter oligomers dominate the degradation profile of stereocomplex PLA, which our data corroborates. Going further, we examined PDLA extracts and observed that PDLA degrades more readily than either PLLA or stereocomplex PLA, consistent with in-vivo degradation profiles in humans^1^.

Making and characterizing PLA extracts is a standardized process as outlined by the International Standard Organization (ISO 10993-5:2009 - Biological evaluation of medical devices). Over the 12-day extraction for this study, there were no pH changes in serum-containing DMEM medium used. This is likely because DMEM medium is highly buffered with sodium bicarbonate salts for cell culture in CO_2_ incubators, and included serum; within CO_2_ incubators, the pH of DMEM medium normalizes to 7.4^56,57^. Therefore, because pH was similar across groups in our study, acidity was not a confounder in our model, and could not have accounted for the dramatic immune cellular outcomes that we observed. If our extractions are performed in water, we reproduce decreased pH which is consistent with prior in-vitro models because water lacks buffers, such as serum and bicarbonates which are present in blood. Remarkably, stereocomplex PLA extracts in water were more acidic than either PLLA or PDLA extracts as previously reported^4^.

Monomeric lactic acid acts as a potent signaling molecule, transcending its role as simply a metabolite both in inflammation and cancer biology^58,59^. In cancer biology, lactate plays as immunomodulatory role^60^. However, combined with bacterial lipopolysaccharide (LPS), lactate could stimulate^51^ or dampen^61^ the activation of immune cells.

For many types of polylactide, reprocessing (by melt-blending) results in faster degradation^62^ which could accentuate immune cellular responses. However, that is not the case with stereocomplex PLA which has been reported to degrade slower than either homopolymer despite (owing to) melt-blending^4,62,63^. Consequently, whereas we expected lower immune cellular activation from stereocomplex PLA since it breaks down in a particularly slow manner, we observed quite the opposite.

Intriguingly, PLLA or PDLA degradation products increased MCP-1 (CCL-2) protein levels whereas stereocomplex PLA did not. MCP-1 recruits macrophages to promote inflammation. MCP-1 is also implicated in cartilage destruction and osteolysis, and is required for foreign body giant cell formation ^64–66^, events that characterize the host immune responses to PLLA and PDLA in-vivo. These different immune cellular responses to PLA of varied stereochemistries could explain conflicting in-vivo study results showing mild and severe host immune responses to different types of PLA^7–17^.

Irrespective of the proinflammatory protein (s) generated by the biomaterials studied, targeting metabolism using 3PO, 2DG or a.a. decreased undesirably high cytokine protein levels. Concomitantly, this strategy markedly increased anti-inflammatory cytokines which are crucial for tissue repair and regeneration through remodeling and recruitment of stem or progenitor cell populations ^48^. Next generation biomaterials used in diagnostic, therapeutic and regenerative applications will be designed to allow for modulation of their immune microenvironment^1^. Targeting bioenergetics and altered metabolism in immune cells offers opportunities for modulating immune responses for improved outcome. Manipulating immune cell activation using biologically-derived, decellularized extracellular matrix is very promising ^67,68^ but complete removal of cellular components, which elicit rejection and risk disease transmission, remain limiting factors^69^; methods aimed at bioenergetics and altered metabolism overcome these limitations.

This study is not focused on the type of PLA that minimizes adverse immune responses but investigates the complex relationship between PLA stereochemistry and immunometabolism. Different types of PLA may be appropriate for different clinical situations, based on their physicochemical properties. For example, highly crystalline PLA with high mechanical strength could be desirable for implants expected to degrade slowly following surgical implantation; whereas, amorphous PLA with reduced mechanical strength could be applied in scenarios where fast degradation is expected. PLA stereochemistry is a key determinant of several of these physicochemical parameters affecting clinical use. By targeting metabolism, we can modulate immune responses to PLA of varied stereochemistries. This could significantly extend the clinical applications of PLA past PLLA and PDLA to formulations of different stereochemistries with minimal concerns about biocompatibility.

The biocompatibility of PLA degradation products presents with conflicting events, partly because “polylactide (PLA)” refers to a broad class of polymer having vast physicochemical, mechanical and thermal properties which are all affected by its stereochemistry. Our findings provide insight into these discrepancies by demonstrating fundamentally different bioenergetic and metabolic signatures in immune cells exposed to degradation products of PLLA, PDLA and their 50/50 combination as stereocomplex PLA. Consequently, varied macrophage polarization occurs. This drives distinctive cytokine and chemokine secretion, ultimately determining host immune responses. In addition, we present a unifying mechanism by which different forms of PLA drive adverse host responses and new methods to specifically modulate them. Except for 3PO, using glycolytic inhibitors allow for some proinflammatory cytokines to be produced by macrophages. This is important because an appropriate level of inflammation is crucial for tissue healing and regeneration to occur. Moreover, the multiple strategies we have used to modulate immune cellular responses offer options from which to choose and apply in a patient-specific manner.

## 5. Conclusion

While different immune outcomes occur following surgical implantation of various types of PLA^7–17^, this observation has not been previously modeled in-vitro. Not only do we model this phenomenon, we reveal the biological mechanisms that explain the complex relationships among material stereochemistry, immune responses, bioenergetics and metabolic reprogramming. Our findings underscore PLA stereochemistry as a determinant of immune cellular responses. With stereocomplex PLA, expression of proinflammatory cytokines is mechanistically driven by increased glycolytic flux in macrophages. Whereas both oxidative phosphorylation and glycolysis are increased with PDLA, only oxidative phosphorylation is increased with PLLA. Taken together, we: 1) highlight the intricacies underlying metabolic alterations among different cell populations in the immune microenvironment after exposure to PLA of varied stereochemistries; 2) offer mechanistic insights into why various types of PLA elicit markedly different immune responses in patients; 3) underscore metabolic reprogramming and altered bioenergetics in immune cells as a unifying mechanism for PLA-induced host responses; and 4) demonstrate immunometabolism as a pivot in biomaterial biocompatibility for controlling host immune responses.

## Author contributions

Conceptualization, C.V.M. and C.H.C.; Methodology, C.V.M., K.D.H., S.B.G., R.N. and C.H.C.; Investigation, C.V.M., M.A., E.U., M.O.B. and M.M.K.; Writing – Original Draft, C.V.M.; Writing – Review & Editing, C.V.M., M.A., E.U., M.O.B., M.M.K., K.D.H., S.B.G., R.N. and C.H.C.; Funding Acquisition, C.H.C.; Resources, R.N. and C.H.C.; Supervision, K.D.H., S.B.G., R.N. and C.H.C.

## Data availability

All data generated during this study are included in this published article and supplementary information files.

## Acknowledgements

AV Makela provided expertise for running cytokine and chemokine assays. Euthanized C57BL/6J mice were a gift from RR Neubig (facilitated by J Leipprandt and E Lisabeth) and the Campus Animal Resources at Michigan State University (MSU). Funding for this work was provided in part by the James and Kathleen Cornelius Endowment at MSU.

## Appendix A. Supplementary data

Supplementary data to this article can be found online at…

