## Supplementary Information for "Stereochemistry Determines Immune Cellular Responses to Polylactide Implants"

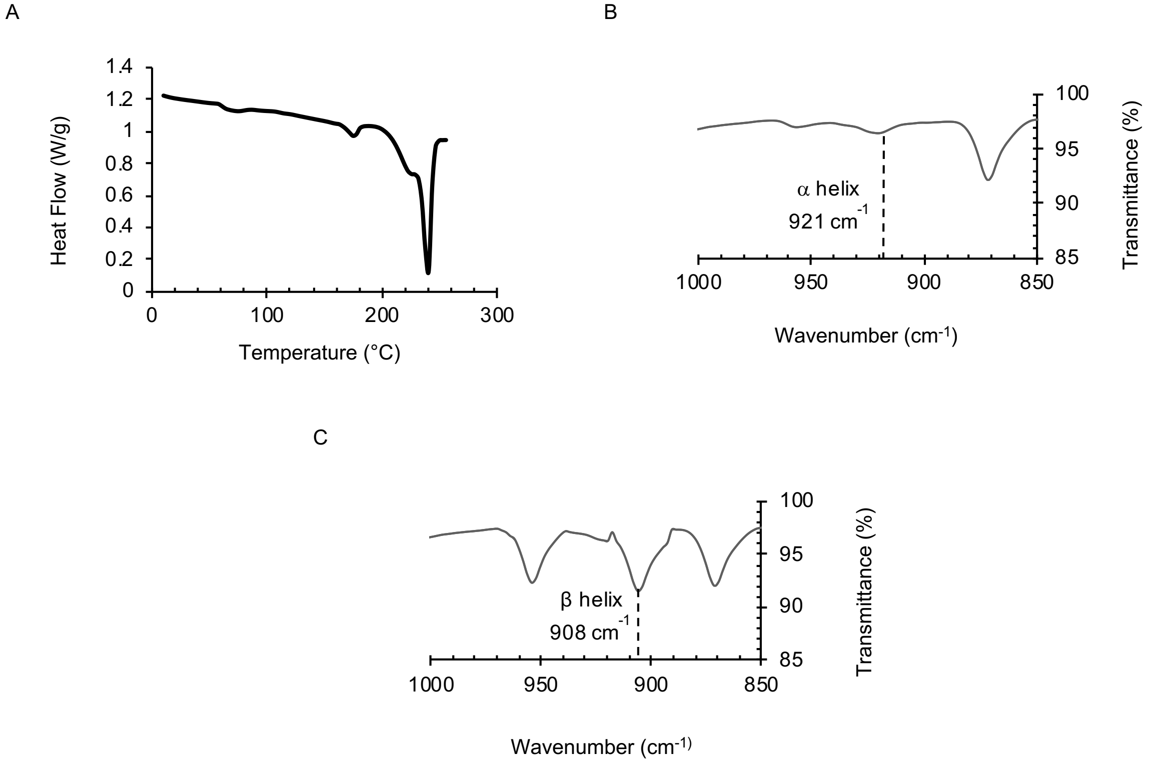


**Fig. S1**. A) Differential scanning calorimetry (DSC) thermogram for the first heating scan of stereocomplex polylactide (PLA) suggests stereocomplexation occurs. B-C) Attenuated total reflectance–Fourier transform infrared (ATR–FTIR) spectroscopy of PLLA (B) and stereocomplex PLA (C) shows their characteristic helices.

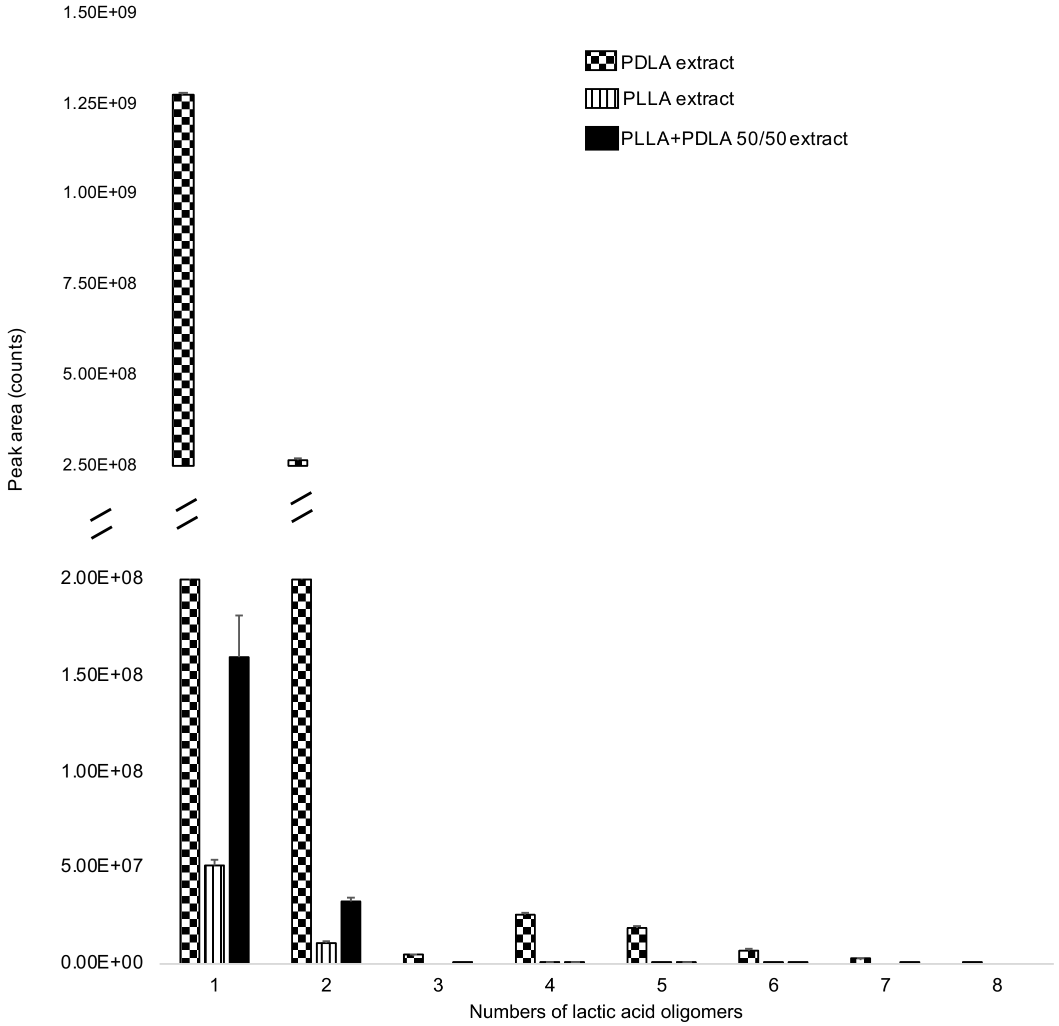


**Fig. S2.** Peak areas of lactic acid oligomers obtained by liquid chromatography-electrospray ionization mass spectrometry of milliQ water-derived extracts of polylactide containing >99% L-isomer (PLLA), >99% D-isomer (PDLA) or a 50/50 melt-blend of PLLA and PDLA (stereocomplex PLA); mean (SD) of n = 3 samples.

5

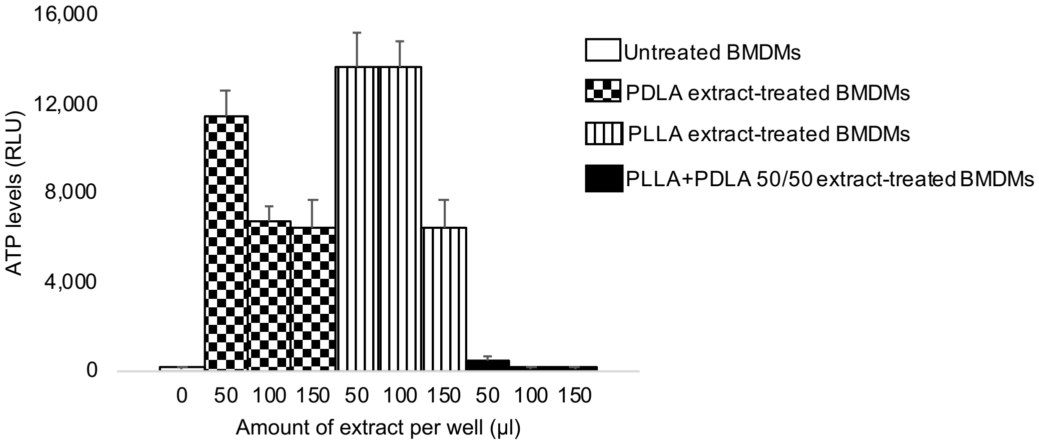


**Fig. S3**. Dose-bioenergetic response of >99% L-isomer (PLLA), >99% D-isomer (PDLA) or a 50/50 melt-blend of PLLA and PDLA (stereocomplex PLA) extracts on primary bone marrow-derived macrophages (BMDMs) reveals an inverse relationship, and tendencies to alter ATP levels for all tested doses. Mean (SD), n = 3, measurements were obtained on day 7.

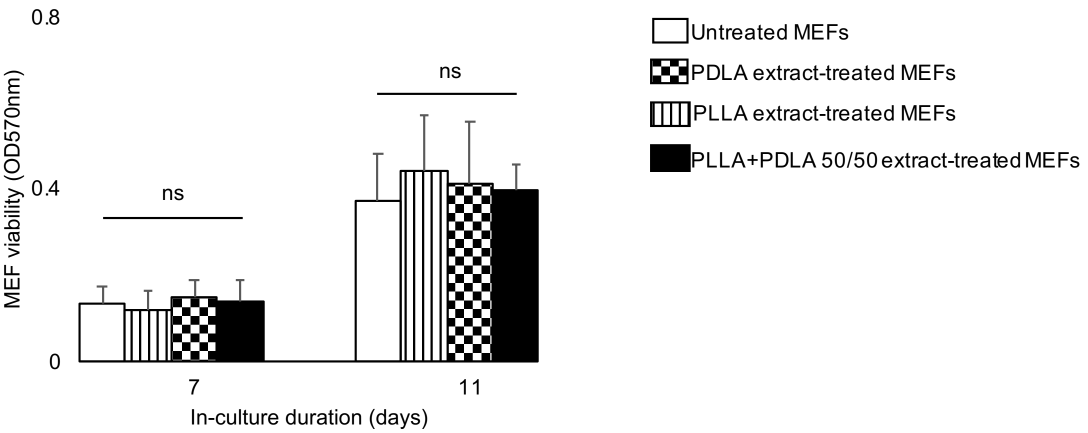


**Fig. S4**. Cell numbers are similar after exposure of mouse embryonic fibroblasts (MEFs) to >99% L-isomer (PLLA), >99% D-isomer (PDLA) or a 50/50 melt-blend of PLLA and PDLA (stereocomplex PLA) extracts over time. Not significant (ns), mean (SD), n = 5, one-way ANOVA; 150 µl of control or extract was used.

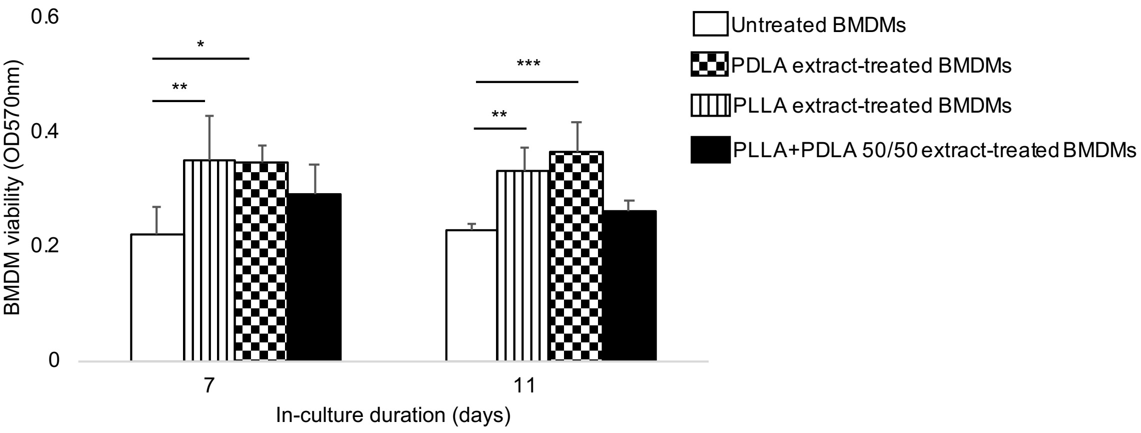


**Fig. S5**. Numbers of primary bone marrow-derived macrophages (BMDMs) are higher after exposure to >99% L-isomer (PLLA) and >99% D-isomer (PDLA) and not a 50/50 melt-blend of PLLA and PDLA (stereocomplex PLA) extracts over time. *$p<0.05$, **$p<0.01$, ***$p<0.001$, mean (SD), n = 5, one-way ANOVA followed by Tukey's post-hoc test; 150 µl of control or extract was used.

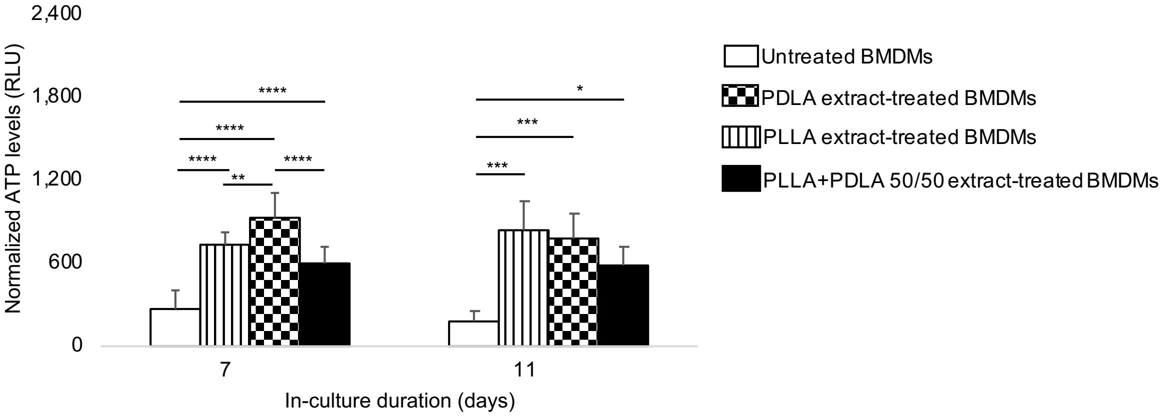


**Fig. S6**. After normalization to cell numbers, bioenergetics in primary bone marrow-derived macrophages (BMDMs) remains altered after prolonged exposure to extracts of polylactide containing >99% L-isomer (PLLA), >99% D-isomer (PDLA) or a 50/50 melt-blend of PLLA and PDLA (stereocomplex PLA) over time. *p<0.05, **p<0.01, ***p<0.001, ****p<0.0001, mean (SD), n= 4-10, one-way ANOVA followed by Tukey's post-hoc test; 150 µl of control or extract was used; normalization factors were obtained from Figure S3 as 1.6, 1.6 and 1.3 for PLLA, PDLA and stereocomplex PLA, respectively, on day 7; 1.4, 1.6 and 1.4 for PLLA, PDLA and stereocomplex PLA, respectively, on day 11.

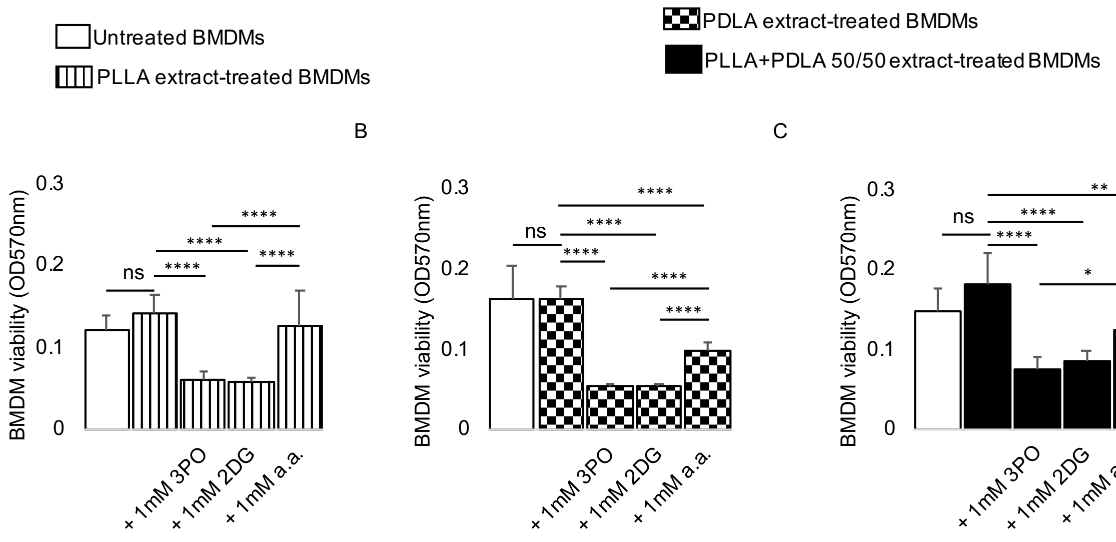


**Fig. S7**. A-C) In comparison to untreated cells, viability of primary bone marrow-derived macrophages (BMDMs) is similar after exposure to polylactide containing >99% L-isomer (PLLA) or >99% D-isomer (PDLA) or a 50/50 melt-blend of PLLA and PDLA (stereocomplex PLA); addition of glycolytic inhibitors reduces cell viability, with aminooxyacetic acid (a.a.) having the least effect. Not significant (ns), *$p<0.05$, **$p<0.01$, ****$p<0.0001$, mean (SD), n = 8, one-way ANOVA followed by Tukey's post-hoc test or Brown-Forsythe and Welch ANOVA followed by Dunnett multiple comparison test; 3-(3-pyridinyl)-1-(4-pyridinyl)-2-propen-1-one (3PO), 2-deoxyglucose (2DG); 100 µl of control or extract was used for 7 days.

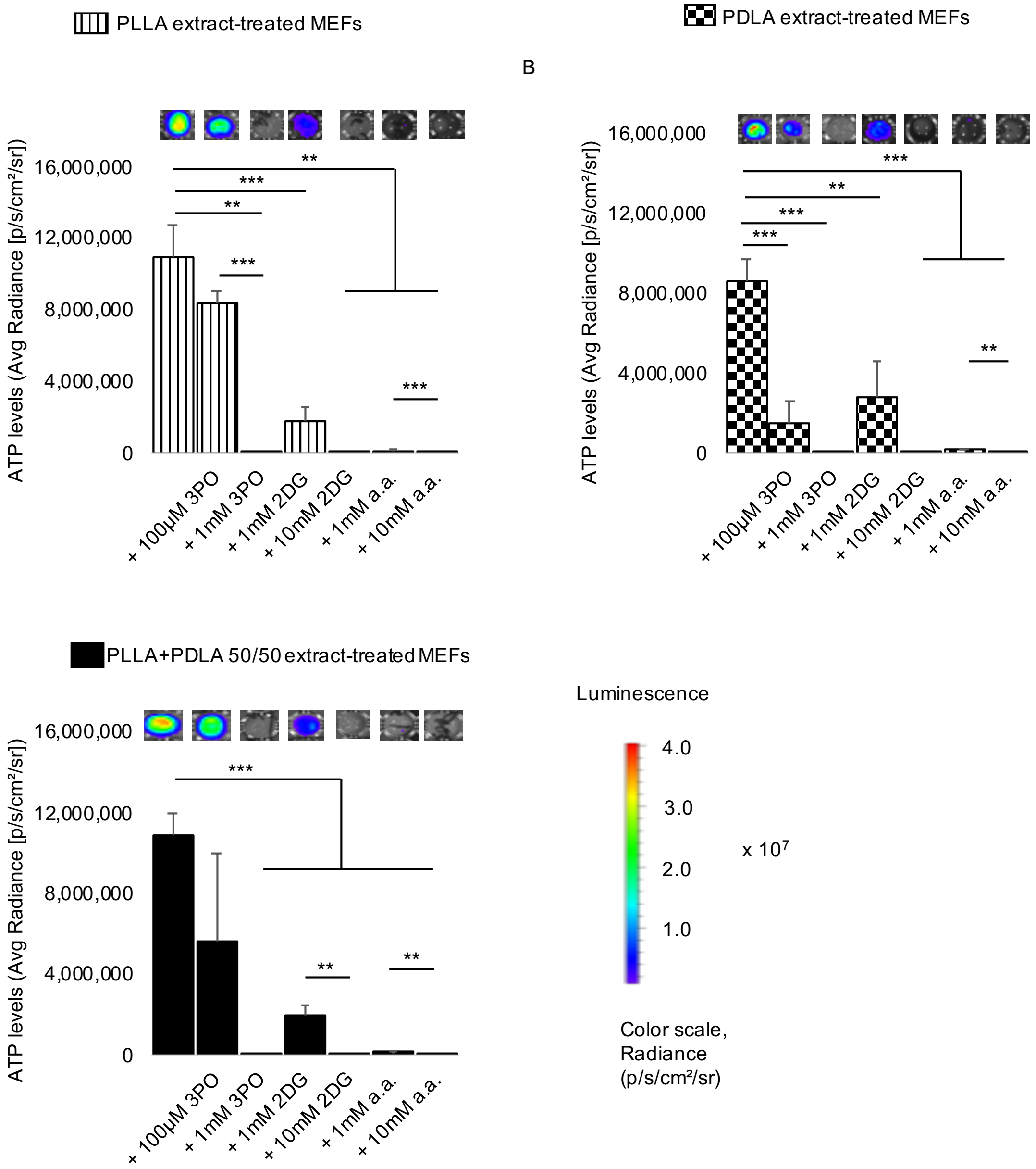


**Fig. S8**. A-C) Bioenergetics is modulated in mouse embryonic fibroblasts (MEFs) exposed to extracts of polylactide containing >99% L-isomer (PLLA) or >99% D-isomer (PDLA) and a 50/50 melt-blend of PLLA and PDLA (stereocomplex PLA) in a dose-dependent manner by pharmacologic inhibitors of glycolysis (representative wells are shown). $^{**}p<0.01$, $^{***}p<0.001$, $^{****}p<0.0001$, mean (SD), n = 5, Brown-Forsythe and Welch ANOVA followed by Dunnett multiple comparison test; 3-(3-pyridinyl)-1-(4-pyridinyl)-2-propen-1-one (3PO), 2-deoxyglucose (2DG) and aminooxyacetic acid (a.a.); 100 µl of control or extract was used for 7 days.

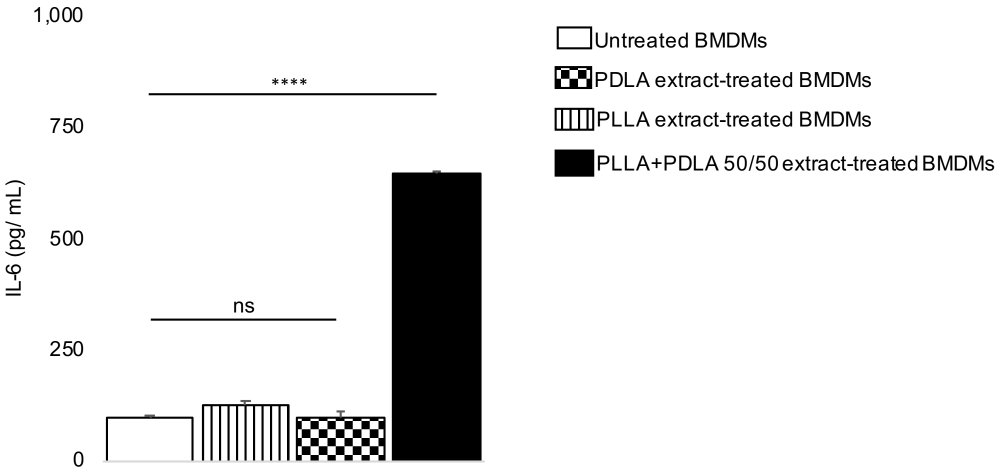


**Fig. S9**. Using ELISA, IL-6 protein expression is similar in primary bone marrow-derived macrophages (BMDMs) exposed to extracts of polylactide containing >99% L-isomer (PLLA) or >99% D-isomer (PDLA) in comparison to untreated macrophages; but is increased when macrophages are exposed to a 50/50 melt-blend of PLLA and PDLA (stereocomplex PLA) extracts. Not significant (ns), ****$p<0.0001$, mean (SD), n = 3, one-way ANOVA followed by Tukey's post-hoc test; 150 µl of control or extract was used for 7 days.

**Table S1.** Physical, chemical and thermal properties of polylactides studied.

| Criteria | PLA L175 (PLLA) | PLA D120 (PDLA) | Stereocomplex PLA (50% PLLA + 50% PDLA) |
|---|---|---|---|
| Optical purity (%) | 99.87 | 99.55 | Not applicable |
| L-content (%) | 99.74 | Not applicable | 50 |
| D-content (%) | Not applicable | 99.87 | 50 |
| Glass transition temperature $T_g$ (ºC) | 63.12 | 62.19 | 64.24 |
| Melting temperature $T_m$ (ºC) | 175.12 | 177.53 | 240.13 |
| Crystallinity (First heating scan, %) | 47.49 | 51.38 | 55.03 |
| Crystallinity (Second heating scan, %) | 6.14 | 36.02 | Not applicable |
| Number average molecular weights $M_n$ (Da) | 102,697 | 91,760 | Not applicable |
| Weight average molecular weights $M_w$ (Da) | 171,675 | 150,515 | Not applicable |
| Polydispersity index | 1.672 | 1.640 | Not applicable |

First heating scan was used to determine $T_m$ and $T_g$ for stereocomplex PLA.
Second heating scan was used to determine $T_m$ and $T_g$ for PLLA and PDLA.
Molecular weights were based on a calibration curve of polystyrene standards.
